# PlotS: web-based application for data visualization and analysis

**DOI:** 10.1101/2023.06.09.544161

**Authors:** Ringyao Jajo, Shivani Kansal, Sonia Balyan, Saurabh Raghuvanshi

## Abstract

**Background:** Data visualisation technique has greatly improved as technology has advanced. While representing the data through graph, it has made the underlying data structure become more transparent and interpretable. However, the informational scope of freely available generic visualisation tools is still limited since they only support descriptive statistics as well as no metainformation can be incorporated. Additionally, there is a need to promote the use of accepted scientific standards when performing and reporting statistical analysis.

**Result:** PlotS is a visualization centric web-based application that allows the integration of statistical analysis into a single workflow. The current version has eight types of graphs (bar, box, density, frequency polygon, histogram, line, scatter and violin plot) and four statistical methods (T-test, ANOVA, Wilcoxon test and Krushkal-Wallis test). It is an interactive application that provide many useful customization options for data analysis-focused visualization. It support incorporation of metainformation in the graph and multivariate data analysis by adding layer with or without secondary y-axis, side graphs, inset graph or through faceting. It can handle variety of data formats with or without replicates and inferential statistical analysis can be incorporated in the graph. Necessary statistical results are explicitly displayed for inferences and reporting.

**Conclusion:** PlotS is freely available at https://plots-application.shinyapps.io/plots/. We hope it will be a useful tool for data visualisation and analysis that will also encourage the widespread use of proper statistical methods in research and teaching.

## Background

Visualization aids in data interpretation and allows scientists to effectively communicate their findings. With the advancement of technology and awareness of computer knowledge, the art of conveying information, data structure, and analysis process through figure has significantly improved. The ongoing effort to maximise the expressivity of figures for improved interpretability has resulted in the development of free and novel visualisation methods ^1,2^. The dynamic nature of the tools allows user to investigate the relationship between variables in different aspect. However, freely available visualisation tools are still limited in the information they convey to the user for interpretation. It does not support incorporation of metadata as well as support only descriptive statistics like measure of dispersion, frequency, etc., which is not adequate for more robust analysis. Therefore, user often relied on alternative statistical tools to conduct inferential statistics. Majority of such tools are of proprietary products and may not be adept as the visualisation tools for generating figure. While programming language like R and python are excellent alternatives, biological researchers may not always be familiar with or be comfortable to learn the languages. Therefore, an application that supports interactive visualization features with the capability to incorporate inferential statistics in real-time will leverage the analytics potential. It will enable hypothesis testing, convey more information, and represent the finding by producing publication-quality figures in a single workflow without interfering the user’s ease of data analysis. Furthermore, statistical report, particularly in the life sciences, frequently fail to adhere to accepted scientific standards ^3,4^. Moreover, researchers often fail to include effect size in the report either due to lack of awareness or knowledge, or because of inexperience with software ^5^. Effect size is an important statistical outcome for inferences (**Figure**) ^4–6^. For transparency and credibility, it is always recommended to follow proper reporting standards. Here we described an open-access web-based application called PlotS, that supports incorporation of statistical analysis in data visualization within a single workflow as well as allow multivariate analysis. It is also designed to encourage the use of proper statistical methods and reporting the analysis.

### Implementation

#### Web page overview

The application is written in R programming using shiny, ggplot2, tidyverse, rstatix and other packages and hosted at https://www.shinyapps.io/. PlotS is freely accessible online at https://plots-application.shinyapps.io/plots/. Details of the packages are available in the help section of the web page. The generated graph can be downloaded in different formats such as eps, pdf, png, svg and tiff. User can set the image resolution and dimension for downloading the graph. After the end of each session, all data are erased.

PlotS web page is divided into main section and sub-sections. Currently there are three main sections – *About, Visualize* & *analyze* and *Help*. The **About** section provide a brief overview of the application and *Help* section provides quick start guides and example. The *Visualize & analyze* is the section where all data import and analysis are conducted. This section is further divided into three sub-sections – *Data, Graph* and *Summary* (**Figure 1**). The *Data* section provides option to upload file or choose example dataset and manage data. All features related to visualization as well as parameters require for performing statistical analysis are provided in the *Graph* section. Additionally, this page has options to filter data based on variables. The *Summary* is the section where data summary, descriptive statistics and result for inferential statistics are displayed. It also include some features to tune the parameters for the statistical methods.

**Figure 1.**
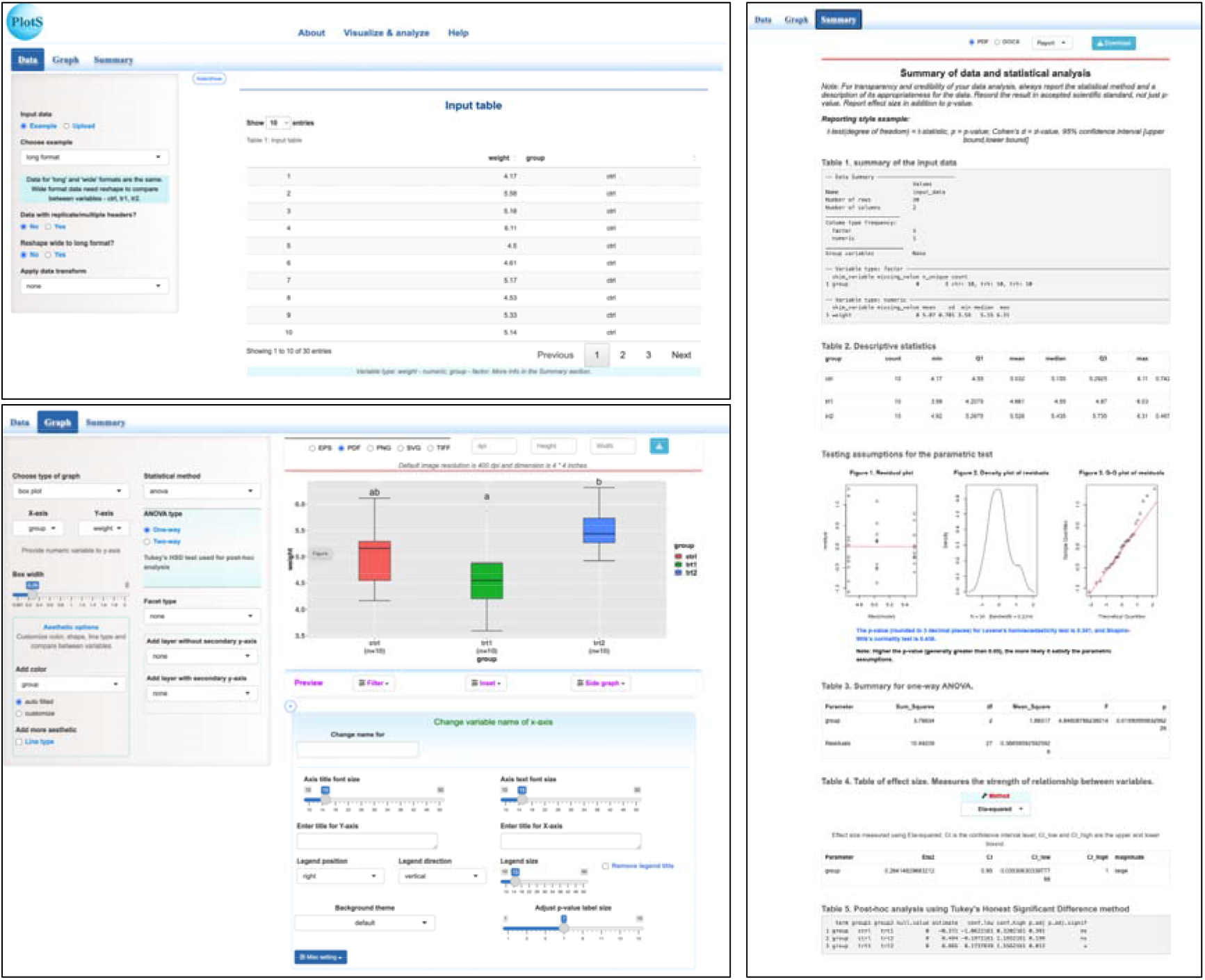
PlotS graphical user interface. Shown only the sub-sections of Visualize & analyze.

### Using the tool

User need to navigate to *Visualize & analyze* section to analyze their data. In the *Data sub-section*, it provides all the utilities useful for data wrangling. User can choose to upload file or use the example dataset. It support variety of file formats such as spreadsheet, comma-separated, tab-separated files, etc for upload. While uploading the file, user can specify multiple missing values. For plotting graph, the app uses the functionality and capability of ggplot2 of R package ^7^. So it provides the flexibility to add multiple variables in the graph to compare interactively during the analysis. Therefore, data can be a mix of non-numeric and numeric columns. Depending on the type of visualisation required for the analysis, it can also be in long or wide formats. However, in most cases, it is necessary to be in long format so that variables can be compared, and the wide format can be easily reshaped to long format using the *reshape* option. It is also designed to handle data with replicates. Replicate data is expected to have more than one header. PlotS allows users to directly import table with multiple headers and format it. The replicates can be pooled and their mean or median determined. The data can also be transformed using different methods. Methods available for transformation are common and binary logarithm, square-root, box-cox and scale. Scale is the function available in R-base. It subtracted the column mean from each row and divided the (centered) column by the standard deviation. The processed data can be downloaded as comma separated file.

Once the data is imported and ready for analysis, it has to navigate to *Graph* sub-section. The current version offers eight types of graphs (bar, box, density, frequency polygon, histogram, line, scatter and violin plot) and four commonly used statistical methods in biological research (parametric: T-test (Welch’s and Student’s test) and ANOVA (one-way and two-way); non-parametric: Wilcoxon test and Krushkal-Wallis test). This section provides various features for visualization. User can customized color and line type based on variables using the aesthetic options. It can add additional layers like point, line or smooth line. There are four smoothing methods to choose – linear regression model (LM), generalized LM, generalized additive model and local regression (LOESS). In addition to this, user can add another layer with secondary y-axis. The units or scales of primary and secondary y-axes can differ (**Figure 3A**). The graph for the secondary y-axis can be bar, box, line or scatter plot.

A separate panel is provided to add inset, side graph or apply filter to the data. Side graph and inset option can be used to add more metainformation to the main graph using bar, box, density, violin plots, etc. Side graph can be drawn on both sides (x- and y-axes) of the main graph (**Figure 2A**). To add an inset, user need to click and drag the mouse around the data point in the main graph. The information of the selected data points can be displayed using different graphs as an inset to the main graph (**Figure 2B**). It can be further customize by the user to display different information for the chosen variables or groups. In addition to visualization function of inset, it also display table for the selected data points which can be downloaded as comma-separated file. In addition to this features, user can facet the graph based on variable for multivariate analysis. It has two type of faceting *viz*. wrap (**Figure 3A**) and grid (**Figure 3B**). If the data contains two discrete variables in all possible combinations, grid faceting can be useful. User can use the wrap faceting if there is only one variable with many levels.

**Figure 2.**
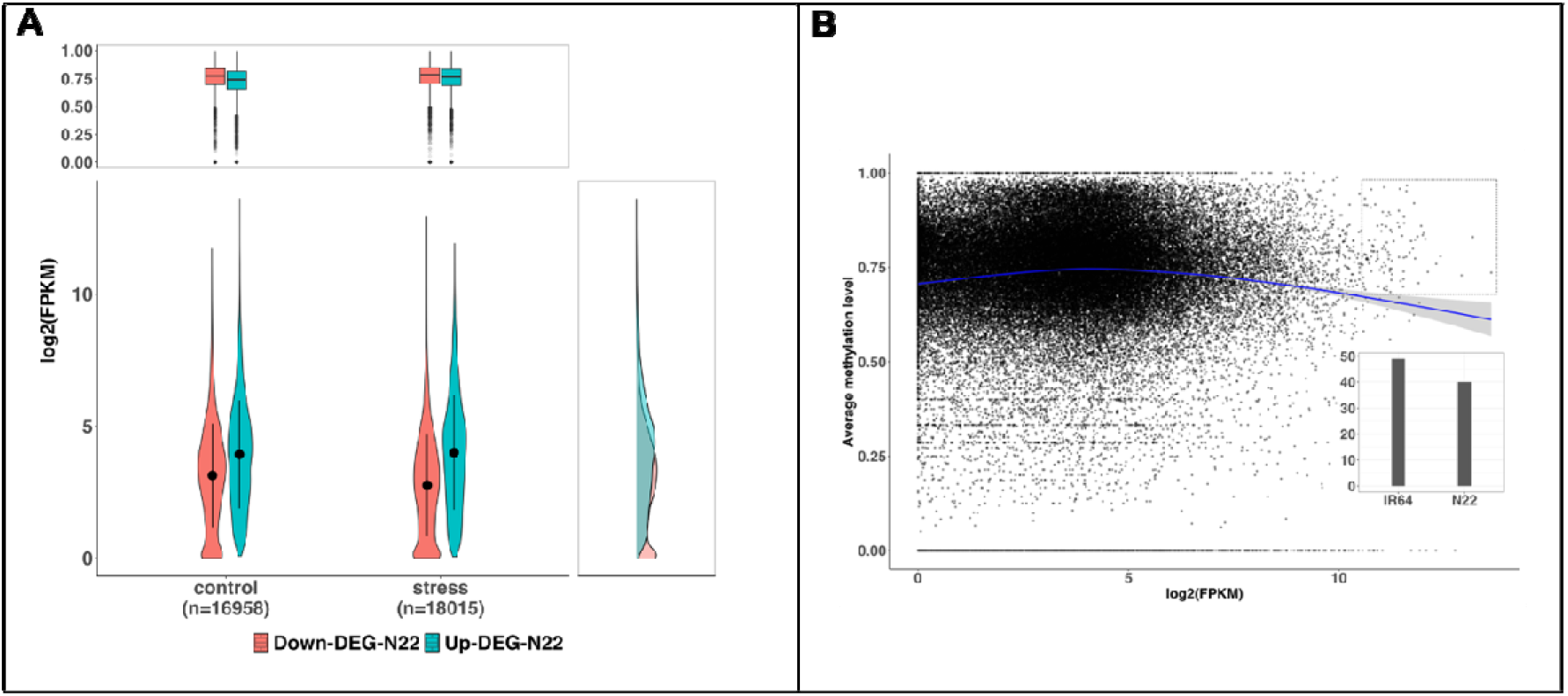
A) **Figure with meta data information**. Main panel shows the transcript level distribution of differentially expressed genes (DEGs) between N22 and IR64 rice cultivars under control and drought stress. The upper panel shows the average methylation level of the DEGs within 1kb upstream to 1kb downstream of transcription start site and the right side panel represents the density of DEGs in the given range of expression under the growth conditions. Point range within the curve represents the standard deviation. “n” in the bracket indicates number of genes. Down-DEG-N22 and Up-DEG-N22 indicated down-regulated and up-regulated in N22 with respect to IR64. B). **Figure with inset**. Showing the relationship between expression and average methylation level of DEGs under drought stress. Blue line represents the linear smooth line (generalized additive model) with confidence interval shaded around the line. Dotted squared area represents the highly expressed DEGs associated with high methylation level. The number of genes for the two cultivars of the selected data are represented in the inset as bar plot.

**Figure 3.**
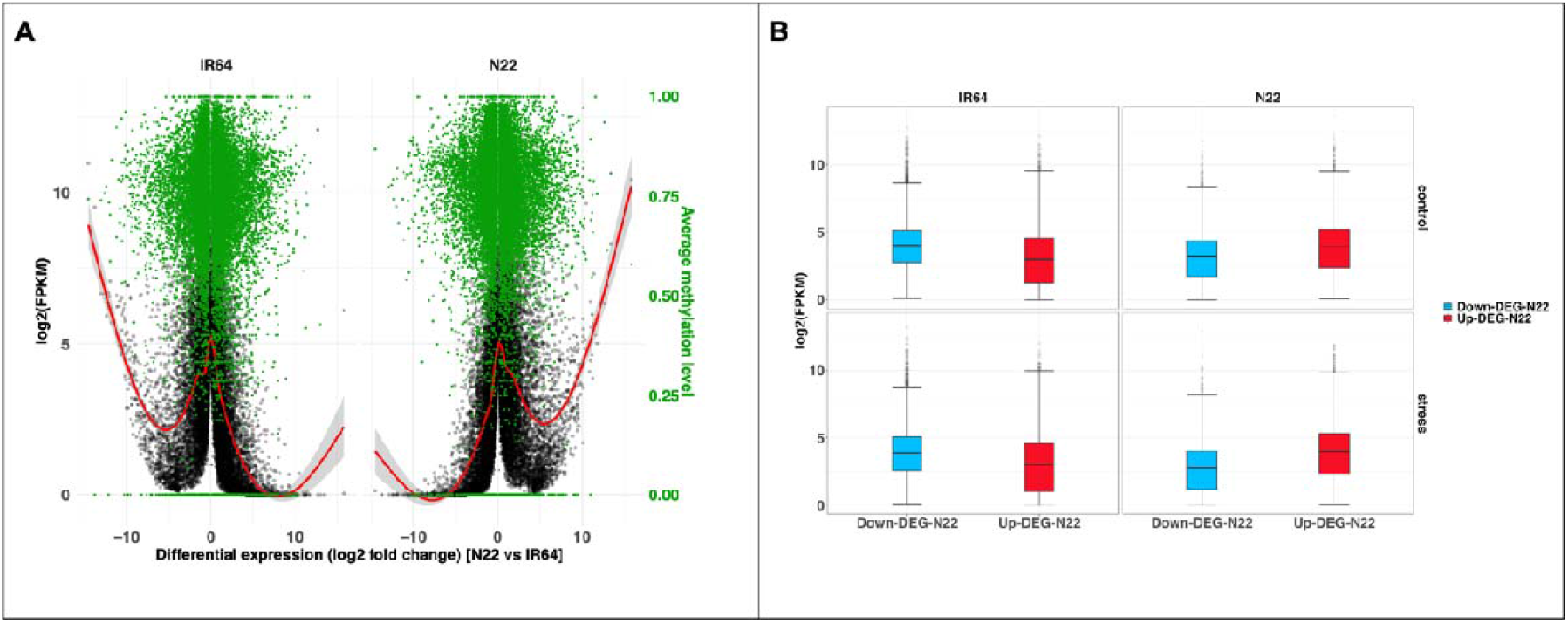
**A)**. Relationship between transcript level, change in expression between cultivars and average methylation level. Differences in the expression of N22 vs IR64 (log2 fold change) is in the x-axis. Positive value indicates higher expression in N22 and negative value means higher in IR64. The primary y-axis is the transcript levels (log2 FPKM) of genes and the average methylation level of genes constitutes the secondary y-axis. The black dots represent the transcript level and green color dots represent the average methylation of the genes. The red color line is the linear smooth line (generalized additive model) of transcript level of genes. Shaded color around the line represents the confidence interval (95%). **B)** Transcript level distribution of differentially expressed genes under control and drought stress in IR64 and N22. Blue color box is for down-regulated genes and red color box represent the up-regulated genes in N22 compared to IR64.

Depending on the type of graph it can also add error bar, a useful feature for describing the data structure and commonly used in research publication ^8^. If the error has been estimated and included in the data then user can specify the column; if not, user need to choose the option to auto compute the error. It has both the features of descriptive (standard deviation) and inferential error bars (standard error and confidence interval [at 95%]). For more robust statistical analysis, user can choose the four statistical methods. Statistical analysis can be performed only after a graph has been chosen. The result will be auto annotated in the graph and the detail of the analysis will be displayed in the *Summary* section.

Statistical methods are implemented using rstatix package and R-base functions. For t-test (Welch’s and Student’s test) and Wilcoxon-test it support both paired and unpaired data. User can also provide group of interest to compare or add group as a reference to conduct the analysis. Two-way ANOVA has feature for both additive and non-additive model. User can also visualized the interaction and main effect of the analysis. For parametric statistics, validity of the tests and effect size are auto generated, so that it is explicitly displayed for inferences. It provides both graphical and statistical test for testing the assumptions. Levene’s test is used for homoscedasticity, and the Shapiro-Wilk’s test is applied for testing normality if the sample size is < 5000; otherwise, the Kolmogorov-Smirnov’s test is used. Statistical result can be downloaded as report or individual table and figure (if applicable).

To estimate the effect size there are different methods for parametric methods which are implemented using rstatix package. For t-test, the methods available are Cohen’s d, Hedge’s g and Glass delta. For ANOVA user can choose Eta-squared, Omega-squared, Epsilon-squared and Cohen’s f. There are two additional methods available for two-way ANOVA *viz*. Partial Eta-squared and Generalized Partial eta-squared. For Wilcoxon and Krushkal-Wallis, we used the non-parametric function available in the package. User can specify the bootstrap for determining the confidence interval at 95%. To perform post-hoc analysis, Tukey Honestly Significant and Dunn’s tests are available for ANOVA and Kruskal-Wallis test, respectively.

## Results

For demonstrating the features of PlotS we used the published RNA-seq data ^9^ and unpublished DNA methylation data of rice generated by the same lab. The two dataset were merged into a single file for analysis. The data are included as supplementary materials. Additionally, for illustration purpose of importance of effect size, we generated normally distributed random data using R language.

### Adding metainformation in the graph

User can incorporate useful information and visualize in a single graph using the features for adding metainformation. The RNA-seq data has genes that are differentially expressed (DEG) between the two cultivars (N22 vs IR64) under control and drought stress. By applying filters on this test data we analysed the distribution of transcript level (FPKM) of N22 cultivars and added the methylation information in the upper panel (**Figure 2A)**. As expected, methylation level of up-regulated genes are lower than the down-regulated genes of N22. User can also add further information on the y-side of the graph as shown in the figure depicting the density of genes. Metainformation can also be added using inset during data analysis (**Figure 2B**). This features is particularly useful for highlighting a finding and for exploratory analysis. The dotted squared area in the main figure represents the highly expressed genes associated with high methylation level. The number of genes for IR64 and N22 within the selected data is shown as inset of bar plot in the figure. There were higher number of genes for IR64 compared to N22 in the selected data.

### Visualizing multivariate data

In addition to adding metadata to the graph, user can also visualized and analyse multiple variables using the features for adding layer with secondary y-axis and facet (**Figure 3**). Including multiple variables in one figure may not always be encouraging, but it is indispensable in cases where complex relationship between variables needs to be explained. The **figure 3A** shows that genes with small differences between cultivar and high expression under stress are more likely to have higher variation in methylation level than genes that are differentially expressed between cultivars. Additionally, distribution of transcript level for the up- and down-regulated DEG of the two cultivars under control and drought stress are shown using grid faceting (**Figure 3B**). If the data has more variables it can be further represented in the graph using aesthetic options. Thus, PlotS provides the flexibility to add multiple variables for analysis.

### Importance of effect size

It is important to include the statistical significance and effect size in interpreting and reporting data analysis. In PlotS, statistical significance can be annotated in the graph as p-value or in symbol. In this illustration we use normally distributed random samples, *sample 1* and *sample 2* with mean 200 (SD ± 10) and 199 (SD ± 5) respectively, were analysed using Welch’s t-test. The differences between their mean at 10000 sample size each were statistically significant, *t*_*welch*_ = 9.0, p < 0.0001 (**Figure 4A**), but the magnitude of the differences was negligible as indicated by the effect size (Cohen’s d = 0.13, 95% CI [0.10, 0.15]). However, if the sample size is reduced to 700 each, the differences in the mean of the two samples are no more statistically significant (*t*_*welch*_ = 0.80, p = 0.42) (**Figure 4B**) and effect size remain negligible (Cohen’s d = 0.04, 95% CI [-0.06, 0.15]). Statistical p-value, which indicates the probability of observed differences between groups due to chance is dependent on the sample size, but effect size which communicates the practical significance is independent of sample size. Therefore inferences should be based on both the analysis.

**Figure 4.**
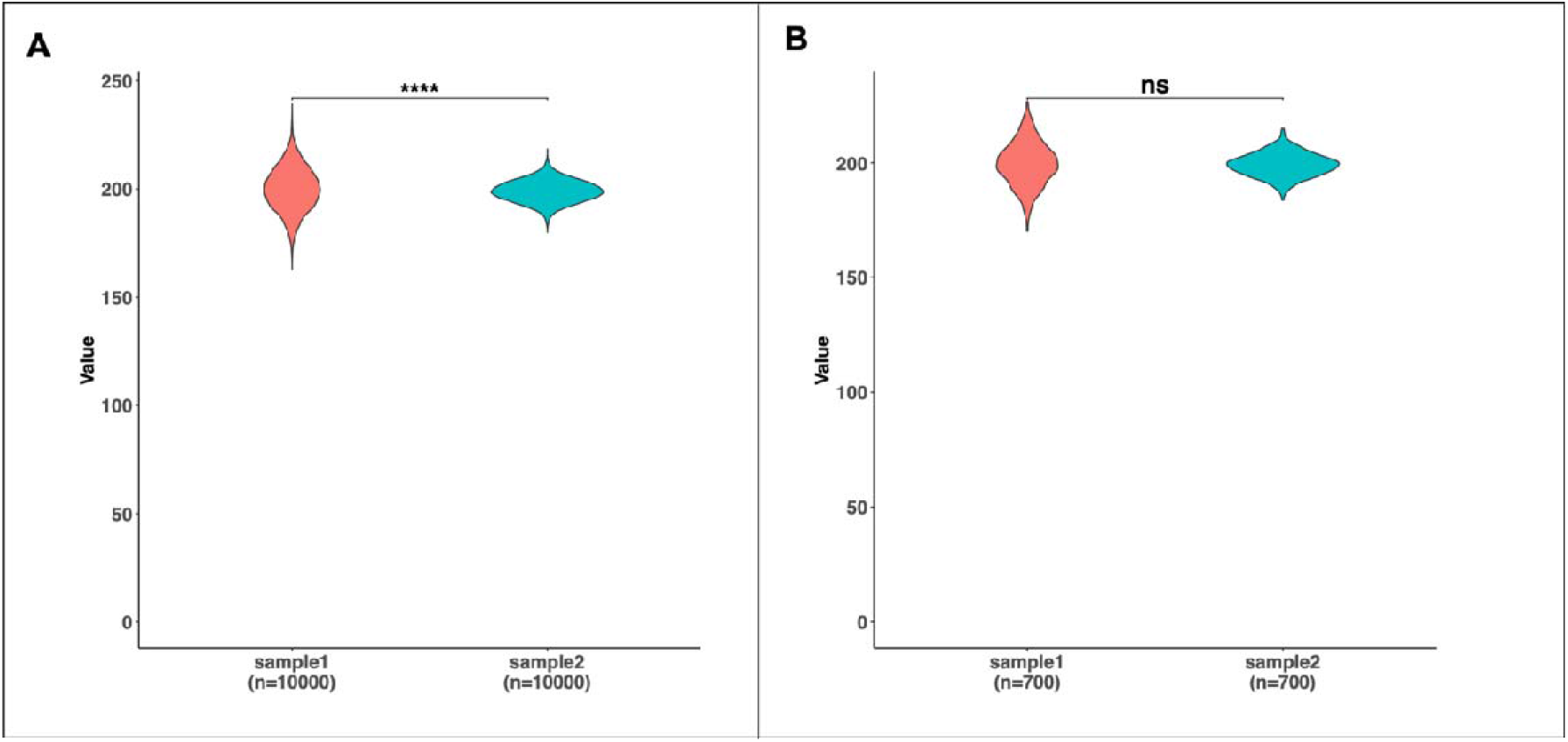
**Statistical annotation in the graph**. Statistical result of t-test (Welch’s test) is annotated as symbol in the graph **A)**. Analysis for 10,000 sample size each. **B)**. analysis for 700 sample size each. Four asterisks indicates p-value < 0.0001 and ‘ns’ indicates no significant differences between the sample mean.

### Limitation

The tool, at present, is unable to handle large data. Another limitation is that PlotS is a data visualization centric app so it does not support standalone statistical analysis, instead all statistical results are generated based on the parameters feed for plotting the graph. To conduct statistical analysis user needs to specify a graph and choose the statistical methods.

### Future update

We are continuing to improve performance and expand visualisation features for heatmaps, upset plots, and cluster analysis. Another significant update we hope to include in PlotS is to add function for recreating an existing graph at any point in time, allowing the user to re-modify the graph or re-analyse the data.

## Conclusion

PlotS allows the integration of statistical analysis and visualization in a single workflow and has the flexibility to include multiple variables in the analysis. It is an interactive web application freely accessible online. It requires no installation.

## Availability and requirements

Project name: PlotS

Project home page: https://plots-application.shinyapps.io/plots/

Operating system: platform independent

Programming language: R

Other requirement: none

License: none

Any restriction to use by non-academics: none

## List of abbreviation

ANOVA: Analysis of Variance
DEG: Differentially Expressed genes
LM: Linear regression Model
LOESS: Locally Estimated Scatterplot Smoothing
SD: Standard Deviation

## Declarations

### Ethics approval and consent to participation

Not applicable

## Consent for publication

Not applicable

## Availability of data and materials

All data generated or analysed during this study are included in this published article

## Authors’ contribution

RJ, SK and SR formulated the concept. RJ wrote the code and drafted the manuscript. SK, SB and SR streamlined the manuscript content. All authors read and approved the final manuscript.

## Acknowledgements

Not applicable

## References

1. Spitzer, M., Wildenhain, J., Rappsilber, J. & Tyers, M. BoxPlotR: a web tool for generation of box plots. Nature Methods 2014 11:2 11, 121–122 (2014).

2. Weissgerber, T. L. et al. Data visualization, bar naked: A free tool for creating interactive graphics. Journal of Biological Chemistry 292, 20592–20598 (2017).

3. Diong, J., Butler, A. A., Gandevia, S. C. & Héroux, M. E. Poor statistical reporting, inadequate data presentation and spin persist despite editorial advice. PLoS One 13, e0202121 (2018).

4. Ordak, M. COVID-19 research: quality of biostatistics. Archives of Medical Science 18, 257–259 (2022).

5. Khalilzadeh, J. & Tasci, A. D. A. Large sample size, significance level, and the effect size: Solutions to perils of using big data for academic research. Tour Manag 62, 89–96 (2017).

6. Ordak, M. Statistical recomendations for the authors of manuscripts submitted to the Journal of Cancer Research and Clinical Oncology. J Cancer Res Clin Oncol 148, 1011–1013 (2022).

7. ggplot2: Elegant Graphics for Data Analysis (3e). https://ggplot2-book.org/.

8. Bella, S., Fidler, F., Williams, J. & Cumming, G. Researchers Misunderstand Confidence Intervals and Standard Error Bars. Psychol Methods 10, 389 (2006).

9. Gour, P. et al. Variety-specific transcript accumulation during reproductive stage in drought-stressed rice. Physiol Plant 174, e13585 (2022).

